# Mechanistic insights into robust cardiac I_Ks_ potassium channel activation by aromatic polyunsaturated fatty acid analogues

**DOI:** 10.1101/2023.01.12.523777

**Authors:** Briana M. Bohannon, Jessica J. Jowais, Leif Nyberg, Sara I. Liin, H. Peter Larsson

## Abstract

Voltage-gated potassium (K_V_) channels are important regulators of cellular excitability and control action potential repolarization in the heart and brain. K_V_ channel mutations lead to disordered cellular excitability. Loss-of-function mutations, for example, result in membrane hyperexcitability, a characteristic of epilepsy and cardiac arrhythmias. Interventions intended to restore K_V_ channel function have strong therapeutic potential in such disorders. Polyunsaturated fatty acids (PUFAs) and PUFA analogues comprise a class of K_V_ channel activators with potential applications in the treatment of arrhythmogenic disorders such as Long QT Syndrome (LQTS). LQTS is caused by a loss-of-function of the cardiac I_Ks_ channel - a tetrameric potassium channel complex formed by K_V_7.1 and associated KCNE1 protein subunits. We have discovered a set of aromatic PUFA analogues that produce robust activation of the cardiac I_Ks_ channel and a unique feature of these PUFA analogues is an aromatic, tyrosine head group. We determine the mechanisms through which tyrosine PUFA analogues exert strong activating effects on the I_Ks_ channel by generating modified aromatic head groups designed to probe cation-pi interactions, hydrogen bonding, and ionic interactions. We found that tyrosine PUFA analogues do not activate the I_Ks_ channel through cation-pi interactions, but instead do so through a combination of hydrogen bonding and ionic interactions.

## Introduction

The delayed rectifier potassium channel (I_Ks_) underlies a critical repolarizing current that determines the timing of the ventricular action potential^1^. The cardiac I_Ks_ current is mediated by the association of the voltage gated K^+^ channel K_V_7.1 α-subunit with the KCNE1 β-subunit^2–4^. The K_V_7.1 α-subunit consists of 6 transmembrane spanning segments, denoted S1-S6 where S1-S4 form the voltage sensing domain (VSD) and S5-S6 form the pore domain (PD)^5^. The S4 segment contains several positively charged arginine residues that allow S4 to move outward, towards the extracellular side of the membrane, when the membrane becomes depolarized^6^. This outward movement of the S4 is transformed into pore opening as a result of conformational changes in the S4-S5 linker of K_V_7.1^7^. Co-expression of KCNE1 with K_V_7.1 imparts a more depolarized voltage-dependence of activation, slower activation kinetics, and increased single channel conductance compared to K_V_7.1 alone^8,9^. Loss-of-function mutations in the cardiac I_Ks_ channel can lead to an arrhythmogenic disorder known as Long QT Syndrome (LQTS), which predisposes individuals to ventricular fibrillation and sudden cardiac death^10–12^. Current treatments for LQTS include pharmacological intervention with β-blockers or surgical implantation of a cardioverter defibrillator^13^. However, limitations of these treatments generate a need for novel therapeutic interventions to treat LQTS.

Polyunsaturated fatty acids (PUFAs) are amphipathic molecules composed of a charged hydrophilic head group and a long, polyunsaturated hydrophobic tail group^14^. It is well-documented that PUFAs form a group of I_Ks_ channel activators that interact with the channel voltage sensing domain (VSD) thus influencing the voltage dependence of I_Ks_ channel activation^15–17^. PUFAs promote I_Ks_ channel activation through an electrostatic interaction between the negative charge of the hydrophilic PUFA head and positively charged arginine residues in the S4 segment of the I_Ks_ channel^17–20^ This electrostatic activation of the I_Ks_ channel is seen as a leftward shift in the voltage dependence of I_Ks_ channel activation that leads to increases in I_Ks_ current. Recently, it has been reported that PUFAs increase I_Ks_ current through two independent effects: one on S4 (as described above) and one on the pore domain through an electrostatic interaction with a positively charged lysine residue located in S6 (K326)^21,22^. This electrostatic interaction with the K326 mediates an increase in the maximal conductance (G_max_) of the I_Ks_ channel^21,22^. The mechanism through which the negatively charged PUFA head group interacts with positive charges of S4 and S6 is called the lipoelectric hypothesis where the polyunsaturated tail of PUFAs and PUFA analogues incorporates into the cell membrane via hydrophobic interactions and electrostatically attracts the outermost gating charges of S4 as well as positively charged K326 in the S6 segment^20,21,23,24^.

PUFA analogues that have the most robust effects on increasing IKs current are those that have a low pKa value and thus possess a negatively charged head group at physiological pH^24^. Examples include PUFAs with glycine or taurine head groups which possess either a carboxyl or sulfonyl head group, respectively^24,25^. We have observed that another PUFA analogue, N-(α-linolenoyl) tyrosine (NALT), has robust effects on I_Ks_ current. NALT is unique in that it possesses a large aromatic tyrosine head group rather than a carboxyl or sulfonyl group present in most of the PUFAs and PUFA analogues that we have characterized. NALT induces a potent leftward shift in the voltage dependence of I_Ks_ channel activation and an increase the maximal channel conductance, thus increasing overall I_Ks_ current. Here, we aim to determine the mechanism behind I_Ks_ activation by NALT using PUFA analogues with aromatic and modified aromatic head groups.

## Materials and Methods

### Molecular Biology

K_V_7.1 and KCNE1 channel cRNA were transcribed using the mMessage mMachine T7 kit (Ambion). 50 ng of cRNA was injected at a 3:1, weight:weight (K_V_7.1:KCNE1) ratio into defolliculated *Xenopus laevis* oocytes (Ecocyte, Austin, TX) for I_Ks_ channel expression. Injected cells were incubated for 72-96 hours in standard ND96 solution (96 mM NaCl, 2 mM KCl, 1 mM MgCl_2_, 1.8 mM CaCl_2_, 5 mM HEPES; pH = 7.5) containing 1 mM pyruvate at 16ºC prior to electrophysiological recordings.

Primers for K_V_7.1 mutations:

R231Q/Q234R – cggccatcaggggTatccAAttTctgAGAatcctgagAatg

K326C – ccagacgtgggtcgggTGCaccatcgcctcctgcttc

S225A – caggtgtttgccacgGCCgcTatcaggggTatccgcttcc

Q220L – gtgggctccaagggAcTTgtgtttgccacgtcgg

T224V – ggggcaggtgtttgcAGTgtcggcTatcaggggcatc

S217A – gtcctctgcgtgggcGccaaggggcaggtgtttg

### PUFA Analogues and Fluorinated PUFAs

Commercially available PUFAs N-(α-linolenoyl) tyrosine (NALT) item number 10032 and linoleoyl phenalanine (Lin-phe) item number 20063 were obtained from Cayman Chemical (Ann Arbor, MI.) or Larodan (Solna, Sweden). Linoleoyl tyrosine (Lin-tyr), Docosahexaenoyl tyrosine (DHA-tyr), and Pinolenoyl tyrosine (Pin-tyr) were synthesized as described previously (Larsson et al., 2020, JGP). NAL-phe, 4Br-NAL-phe, 4F-NAL-phe, 3,4,5F-NAL-phe, and 3F-NALT were synthesized similarly, with detailed descriptions of the synthesis procedures for each compound provided in the supplemental methods. PUFA analogues were kept at −20° C as 100 mM stock solutions in ethanol except 4Br-NAL-phe, 4F-NAL-phe, 3,4,5F-NAL-phe, and 3F-NALT where stock solutions were prepared as needed on the day of recording. Serial dilutions of the different PUFAs were prepared from stocks to make 0.2 μM, 0.7 μM, 2.0 μM, 7.0 μM, and 20 μM concentrations in ND96 solutions (pH = 7.5).

### Two-electrode voltage clamp (TEVC)

*Xenopus laevis* oocytes, co-expressing wild type K_V_7.1 and KCNE1, were recorded in the two-electrode voltage-clamp (TEVC) configuration. Recording pipettes were filled with 3 M KCl. The recording chamber was filled with ND96 (96 mM NaCl, 2 mM KCl, 1 mM MgCl_2_, 1.8 mM CaCl_2_, 5 mM Tricine; pH 9). Dilutions of PUFAs and PUFA analogues were perfused into the recording chamber using the Rainin Dynamax Peristaltic Pump (Model RP-1) (Rainin Instrument Co., Oakland, CA. USA).

Electrophysiological recordings were obtained using Clampex 10.3 software (Axon, pClamp, Molecular Devices). During the application of PUFAs the membrane potential was stepped every 30 sec from −80 mV to 0 mV for 5 seconds before stepping to −40 mV and back to −80 mV to ensure that the PUFA effects on the current at 0 mV reached steady state (Fig. 1D). A voltage-step protocol was used to measure the current vs. voltage (I-V) relationship before PUFA application and after the PUFA effects had reached steady state for each concentration of PUFA. Cells were held at −80 mV followed by a hyperpolarizing prepulse to −140 mV to make sure all channels are fully closed. The voltage was then stepped from −100 to 60 mV (in 20 mV steps) followed by a subsequent voltage step to −20 mV to measure tail currents before returning to the −80 mV holding potential.

**Fig. 1.**
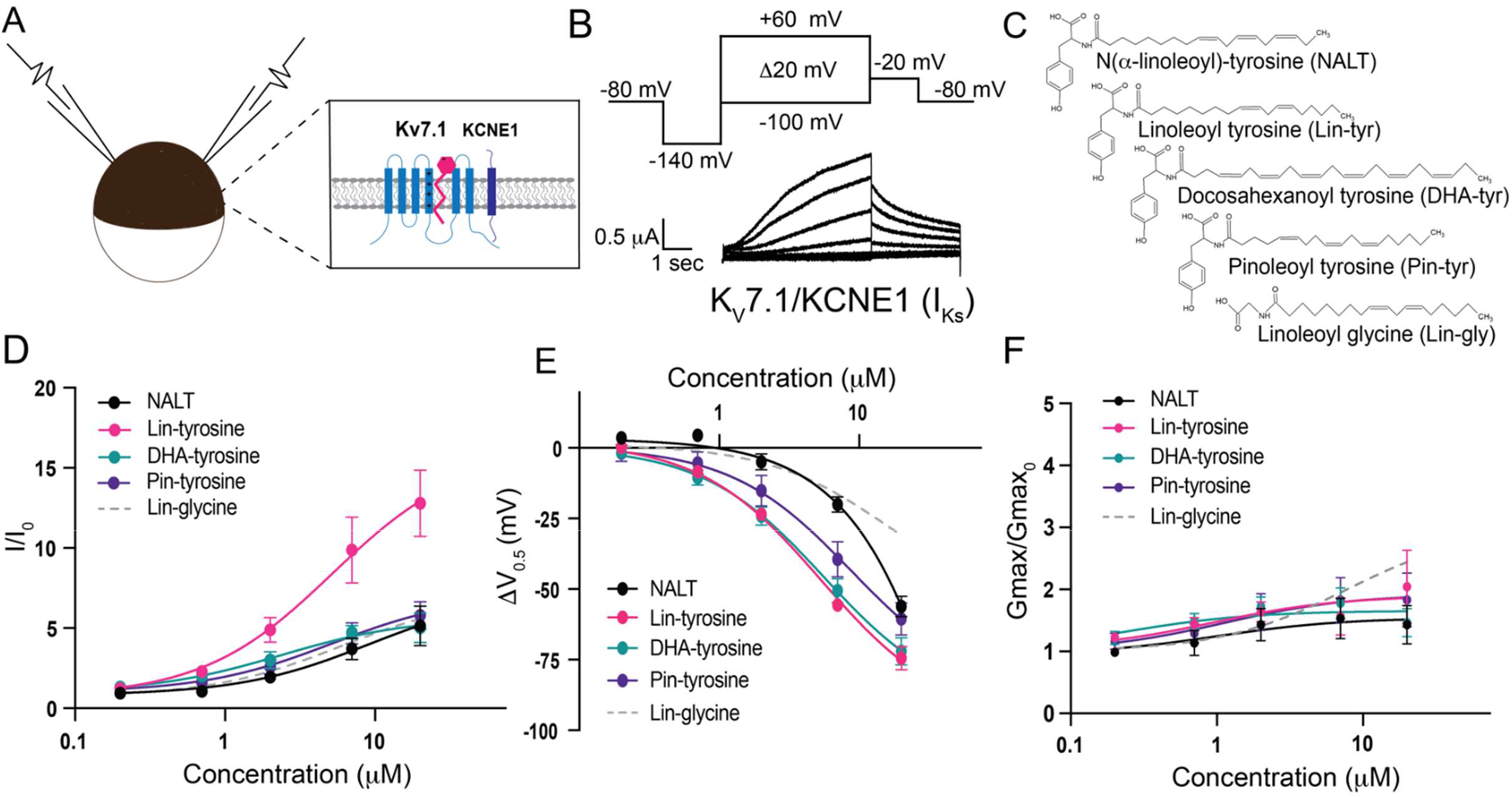
PUFA analogues with a tyrosine head group are strong I_Ks_ channel activators. **A)** Schematic of two electrode voltage-clamp setup (Inset: I_Ks_ channel cartoon + PUFA (pink)). **B)** Voltage protocol (top) with representative K_V_7.1/KCNE1 (I_Ks_) current (bottom). **C)** Structures of NALT, Lin-tyrosine, DHA-tyrosine, and Pin-tyrosine (with Lin-glycine for comparison). **D-F)** I/I_0_, **E)** ΔV_0.5_, and **F)** G_max_ dose response curves for NALT (black circles) (n=4), Lin-tyrosine (pink circles) (n=4), DHA-tyrosine (teal circles) (n=3), Pin-tyrosine (purple circles) (n=5), and Lin-glycine (gray dotted line) (n=3). Values for all compounds and concentrations available in Figure 1-source data 1.

### Data analysis

Tail currents were analyzed using Clampfit 10.3 software in order to obtain conductance vs. voltage (G-V) curves to determine the voltage dependence of channel activation.

The V_0.5_, the voltage at which half the maximal current occurs, was obtained by fitting the G-V curves from each concentration of PUFA with a Boltzmann equation:

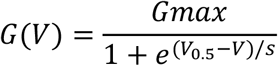

where G_max_ is the maximal conductance at positive voltages and s is the slope factor in mV. The current values for each concentration at 0 mV (I/I_0_) were used to plot the dose response curves for each PUFA. These dose response curves were fit using the Hill equation to obtain the K_m_ value for each PUFA:

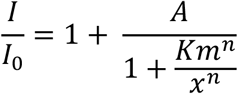

where A is the fold increase in current caused by the PUFA at saturating concentrations, K_m_ is the apparent affinity of the PUFA, x is the concentration, and n is the Hill coefficient. Fitted maximum values derived from the dose response curves are reported for each of the effects (I/I_0_, ΔV_0.5_, and G_max_) from the different PUFAs tested. In some cases, there is variability in the V_0.5_ between batches of oocytes. In order to correct for variability due to oocytes, when the V_0.5_ was greatly different than 20 mV in control solution, we applied a correction in order to more accurately measure PUFA-induced I_Ks_ current increases. We subtracted the V_0.5_ (given by fitting the G-V with a Boltzmann equation) by 20 mV and used the current measured at the resulting voltage. The maximum conductance (G_max_) was calculated by taking the difference between the maximum and minimum current values (using the G-V curve for each concentration) and then normalizing to control solution (0 μM). Graphs plotting mean and standard error of the mean (SEM) for I/I_0_, ΔV_0.5_, G_max_, and K_m_ were generated using GraphPad Prism (GraphPad Software, La Jolla, CA).

### Statistics

Unpaired t-tests and one-way ANOVA with multiple comparisons statistics were computed using GraphPad Prism (GraphPad Software, La Jolla, CA). Results were considered significant if p < 0.05.

## Results

### Diverse PUFA analogues with a tyrosine head group activate the I_Ks_ channel

To measure the effects of the aromatic PUFA analogues on the cardiac I_Ks_ channel, we expressed the I_Ks_ channel complex in *Xenopus laevis* oocytes (Fig. 1A). We co-injected mRNA for the K_V_7.1 α-subunit and the KCNE1 β-subunit to achieve expression of tetrameric I_Ks_ channels. Using two-electrode voltage-clamp recordings, we applied depolarizing voltage steps to activate the I_Ks_ channel (Fig. 1B) before and after applying four different tyrosine PUFA analogues: N(α-linolenoyl)-tyrosine (NALT), Linoleoyl tyrosine (Lin-tyrosine), Docosahexanoyl tyrosine (DHA-tyrosine), and Pinoleoyl tyrosine (Pin-tyrosine) (Fig. 1C). From these voltage-clamp experiments we are also able to acquire dose response curves for different aspects of I_Ks_ channel activation, including changes in overall I_Ks_ current (I/I_0_, Fig. 1D), changes in the voltage-dependence of activation (ΔV_0.5_, Fig. 1E), and changes in the maximal channel conductance (Gmax/Gmax_0_, Fig. 1F). NALT, Lin-tyr, DHA-tyr, and Pin-tyr all activate the cardiac I_Ks_ channel by shifting the voltage dependence of I_Ks_ channel activation to more negative voltages (NALT: −56.2 ± 3.6 mV; Lin-tyr: −74.4 ± 4.1 mV; DHA-tyr: −72.0 ± 4.9 mV; and Pin-tyr: −60.5 ± 5.8 mV at 20 μM; Fig. 1E) and increasing the maximal conductance (NALT: 1.43 ± 0.3; Lin-tyr: 2.0 ± 0.6; DHA-tyr: 1.5 ± 0.2; and Pin-tyr: 1.8 ± 0.4 at 20 μM; Fig. 1F). Together the left shift in V_0.5_ and the increase in G_max_ increase the overall I_Ks_ current measured in response to a voltage step close to 0 mV (NALT: 5.14 ± 1.2; Lin-tyr: 12.8 ± 2.1; DHA-tyr: 5.0 ± 0.9; and Pin-tyr: 5.8 ± 0.9 at 20 μM; Fig. 1D: See Methods for calculation of I/I_0_). In comparison, Lin-glycine (a known I_Ks_ channel activator; Fig. 1C) causes only modest leftward voltage shifts (−30.8 ± 5.4 mV at 20 μM; Fig. 1E), but similar increases in maximal conductance (2.6 ± 0.5 at 20 μM; Fig. 1F) and I/I_0_ current (6.7 ± 1.1 at 20 μM; Fig. 1D) as for most tyrosine PUFAs.

### Distal -OH group is necessary for robust activation of the I_Ks_ channel

Amino acids with aromatic groups (like tryptophan, tyrosine, and phenylalanine) can participate in cation-pi interactions^26^. Cation-pi interactions take place between the pi-electrons of an aromatic ring and positively charged (cationic) groups (such as arginine and lysine)^27^. If tyrosine PUFAs activate the I_Ks_ channel via cation-pi interactions, we would expect that other aromatic groups (such as phenylalanine) would similarly affect I_Ks_ activation. We tested two different PUFA analogues that both contain a phenylalanine head group – Linoleoyl phenylalanine (Lin-phe) and N-(α-linolenoyl) phenylalanine (NAL-phe) (Fig. 2A). Lin-phe and NAL-phe both increase I/I_0_ (Lin phe: 2.6 ± 0.3; and NAL-phe: 2.4 ± 0.5 at 20 μM; Fig. 2B-D), causing a modest leftward shift the V_0.5_ (Lin-phe: −13.1 ± 2.9 mV; and NAL-phe: −12.5 ± 3.8 mV at 20 μM; Fig 2E-F).

**Fig. 2.**
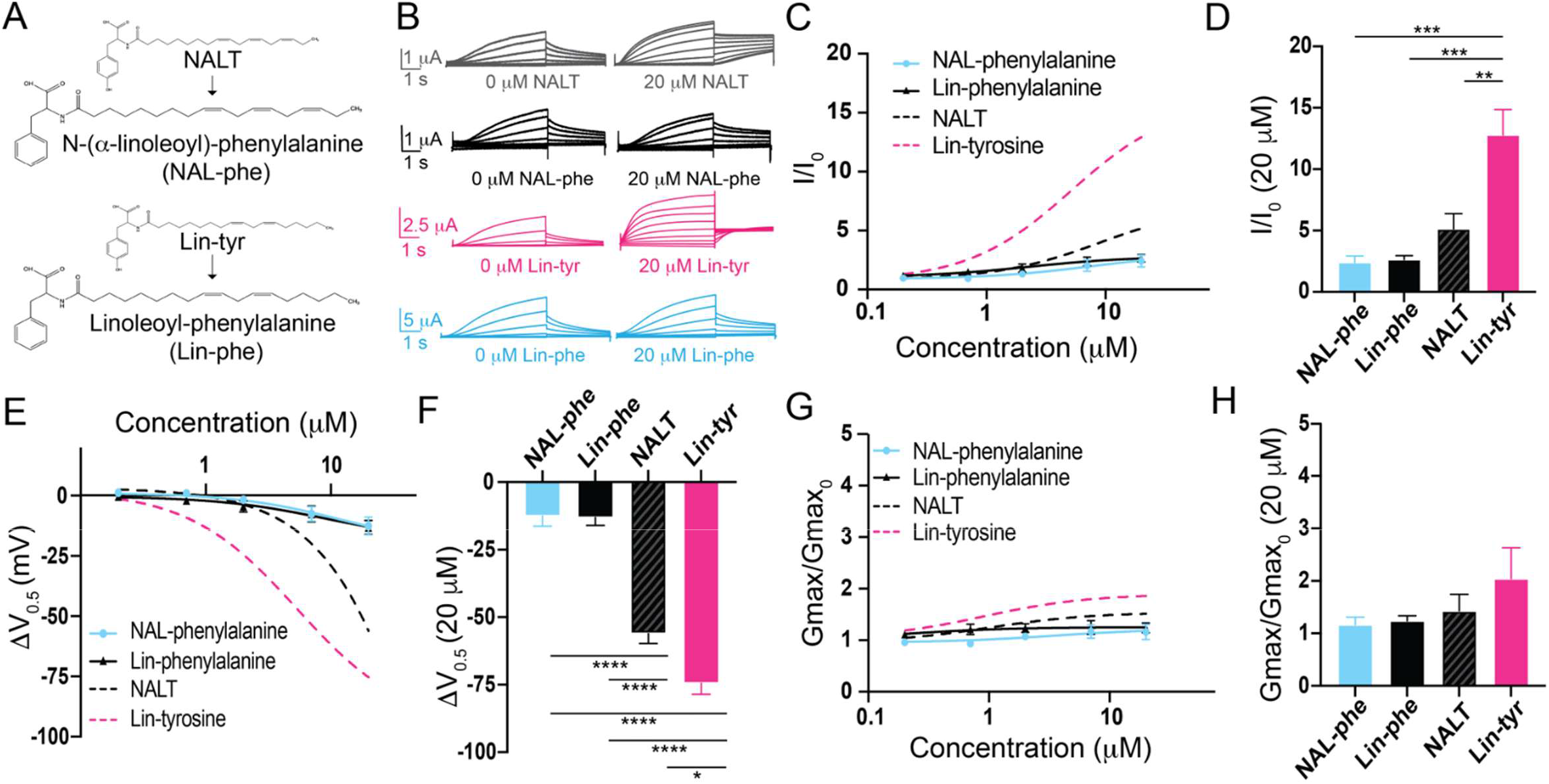
The distal hydroxyl (−OH) group of tyrosine PUFA analogues is necessary for robust I_Ks_ channel activation. **A)** Structures of NAL-phe and Lin-phe. **B)** Representative current traces for NALT (gray), NAL-phe (black), Lin-tyr (pink), and Lin-phe with 0 μM PUFA (left) and 20 μM PUFA (right). **C**,**E**,**G)** I/I_0_, **E)** ΔV_0.5_, and **G)** G_max_ dose response curves for NAL-phe (n=4) and Lin-phe (n=4) with dotted lines representing dose response of NALT (n=4) and Lin-tyr (n=4). **D**,**F**,**H)** Maximum effects on **D)** I/I_0_, **F)** ΔV_0.5_, and **H)** G_max_ (at 20 μM) for NAL-phe (n=4), Lin-phe (n=4), NALT (n=4), and Lin-tyr (n=4). (Asterisks indicate statistically significant differences determined by one-way ANOVA with Tukey’s test for multiple comparisons.) Values for all compounds and concentrations available in Figure 2-source data 2.

However, Lin-phe and NAL-phe have minimal effects on the G_max_ (Lin phe: 1.2 ± 0.1; and NAL-phe: 1.2 ± 0.2 at 20 μM; Fig 2G-H). All of these effects (I/I_0_, ΔV_0.5_, and G_max_) are reduced in comparison with tyrosine PUFAs, with Lin-phe and NAL-phe causing significantly smaller increases in I/I_0_ compared to Lin-tyrosine (p = 0.0004***; Fig. 2D). In addition, both NALT and Lin-tyrosine cause a significantly greater ΔV_0.5_ compared to NAL-phe and Lin-phe (p < 0.0001****; Fig. 2F). Together, these differences suggest that cation-pi interactions are not the primary mechanism through which tyrosine PUFAs activate the I_Ks_ channel. Rather, our data suggest that it is actually the presence of the distal -OH group on the aromatic head group that is critical for the potent activation of the I_Ks_ channel because the loss of this -OH group (Lin-phe and NAL-phe) results in pronounced reductions in PUFA efficacy.

### Electronegative groups on aromatic ring are important for increases in maximal conductance

Our data thus far indicates that it is the presence of the -OH group, not cation-pi interactions, that is critical for pronounced I_Ks_ channel activation by tyrosine PUFAs. The -OH group found in tyrosine PUFAs is highly electronegative. To test how electronegativity influences I_Ks_ channel activation, we compared three modified phenylalanine PUFAs, that all include a highly electronegative group(s) attached to the aromatic ring. We compared N-(α-linolenoyl)-4-bromo-L-phenylalanine (4Br-NAL-phe), N-(α-linolenoyl)-4-fluoro-L-phenylalanine (4F-NAL-phe), and N-(α-linolenoyl) 3,4,5-trifluorophenylalanine (3,4,5F-NAL-phe) (Fig. 3A). 4Br-NAL-phe, 4F-NAL-phe, and 3,4,5F-NAL-phe application increases I/I_0_ (4Br-NAL-phe: 4.6 ± 0.1 at 20 μM; 4F-NAL-phe: 4.6 ± 1.1 at 20 μM; 3,4,5F-NAL-phe: 7.1 ± 1.0 at 20 μM; Fig. 3B-D), causes a leftward shift in the V_0.5_ (4Br-NAL-phe: −22.8 ± 2.0 mV; 4F-NAL-phe: −23.9 ± 0.8 mV; 3,4,5F-NAL-phe: −32.4 ± 4.9 mV at 20 μM; Fig 3E-F), and increases the G_max_ (4Br-NAL-phe: 1.9 ± 0.1 at 20 μM; 4F-NAL-phe: 2.0 ± 0.5; 3,4,5F-NAL-phe: 2.4 ± 0.4 at 20 μM; Fig. 3G-H). Increasing the number of highly electronegative groups significantly improves the effects of phenylalanine PUFAs on increasing I/I_0_ and shifting the V_0.5_, evidenced by significant increases in I/I_0_ (p = 0.0186*; Fig. 3D) and a significantly greater leftward shift in the V_0.5_ (p = 0.0096**; Fig. 3F) from 3,4,5F-NAL-phe compared to NAL-phe alone. Interestingly, though, NALT still causes the most prominent left-shift in the V_0.5_ compared to 4Br-, 4F-, and 3,4,5F-NAL-phe (p = 0.0003***; p = 0.00028***; and p = 0.0021**, respectively). These data suggest that the presence of highly electronegative groups improve the activating effects of phenylalanine PUFAs on the I_Ks_ channel. However, they do not completely recapitulate the effects of tyrosine PUFAs on the shift in V_0.5_ of the I_Ks_ channel.

**Fig. 3.**
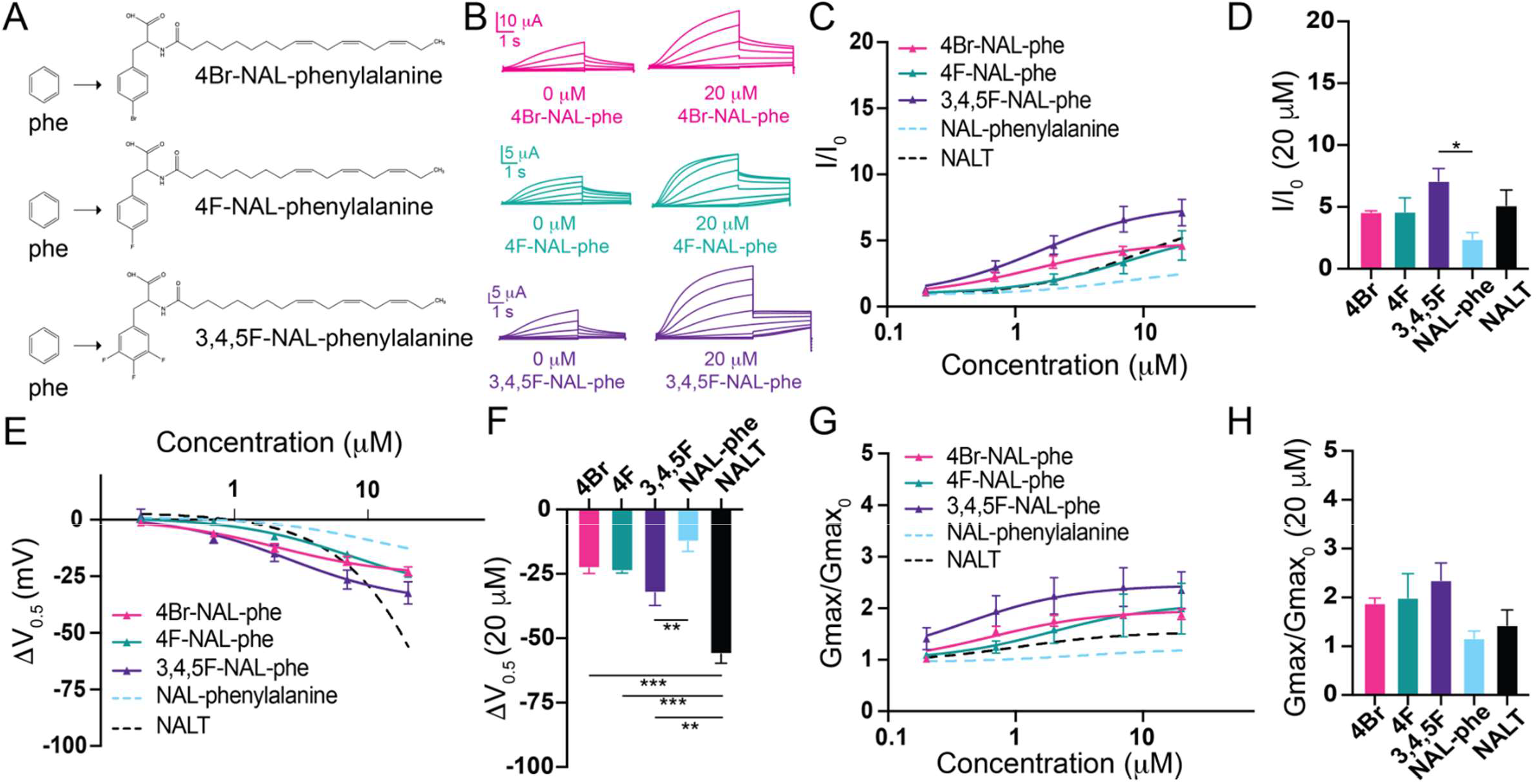
The addition of electronegative atoms to phenylalanine PUFA analogues strengthens I_Ks_ channel activation through improved effects on G_max_. **A)** Structures of 4Br-NAL-phe, 4F-NAL-phe, and 3,4,5F-NAL-phe **B)** Representative traces for 4Br-NAL-phe (pink), 4F-NAL-phe (teal), and 3,4,5F-NAL-phe (purple) with 0 μM PUFA (left) and 20 μM PUFA (right). **C**,**E**,**G)** I/I_0_, **E)** ΔV_0.5_, and **G)** G_max_ dose response curves for NAL-phe (n=4), 4Br-NAL-phe (n=3), 4F-NAL-phe (n=4), and 3,4,5F-NAL-phe (n=5) with dotted line representing dose response of NALT (n=4). **D**,**F**,**H)** Maximum effects on **D)** I/I_0_, **F)** ΔV_0.5_, and **H)** G_max_ (at 20 μM) for 4Br-NAL-phe (n=3), 4F-NAL-phe (n=4), and 3,4,5F-NAL-phe (n=5). (Asterisks indicate statistically significant differences determined by one-way ANOVA with Tukey’s test for multiple comparisons.) Values for all compounds and concentrations available in Figure 3-source data 3.

### Hydrogen bonding is important for pronounced leftward shifts in I_Ks_ channel voltage dependence

The presence of the -OH group on tyrosine PUFA analogues or the addition of electronegative groups to the phenylalanine head group improves I_Ks_ activation. However, a persistent and striking difference between tyrosine PUFAs and modified phenylalanine PUFAs in the magnitude of their voltage-shifting effects with the tyrosine PUFAs having an almost twice as big voltage shift effect than the modified phenylalanine PUFAs (Fig. 3E-F). One explanation for this discrepancy is that the -OH group can also behave as a hydrogen bond donor. To determine if hydrogen bonding contributes to the activating effects of tyrosine PUFA analogues, we applied the modified aromatic PUFA analogue N-(α-linolenoyl)-3-fluoro-L-tyrosine (3F-NALT), which has a fluorine atom adjacent to the tyrosine hydroxyl group (Fig. 4A). The addition of the fluorine atom reduces the pK_a_ of the distal hydroxyl group and increases the hydrogen bonding ability of said group in 3F-NALT as compared to NALT. Overall, the maximum effects on I/I_0_ are similar for 3F-NALT and NALT (3F-NALT: 5.0 ± 1.0; NALT: 5.14 ± 1.2 at 20 μM; p = 0.7257, ns; Fig. 4B-D). Notably, 3F-NALT induces a significantly greater maximum shift in the V_0.5_ (−69.3 ± 1.4 at 20 μM) compared to NALT (−56.1 ± 3.6 AT 20 μM) (p = 0.0298*; Fig. 4E-F), while the effects on G_max_ are not significantly different between 3F-NALT and NALT (3F-NALT: 1.3 ± 0.3; NALT: 1.4 ± 0.3 at 20 μM; p = 0.7324, ns; Fig. 4G-H). These data demonstrate that increasing the hydrogen bonding capacity of the -OH group increases the maximum shift in I_Ks_ channel voltage dependence. This implicates hydrogen bonding as an important mechanism for I_Ks_ activation and preferentially influences the effects on the voltage dependence of I_Ks_ activation.

**Fig. 4.**
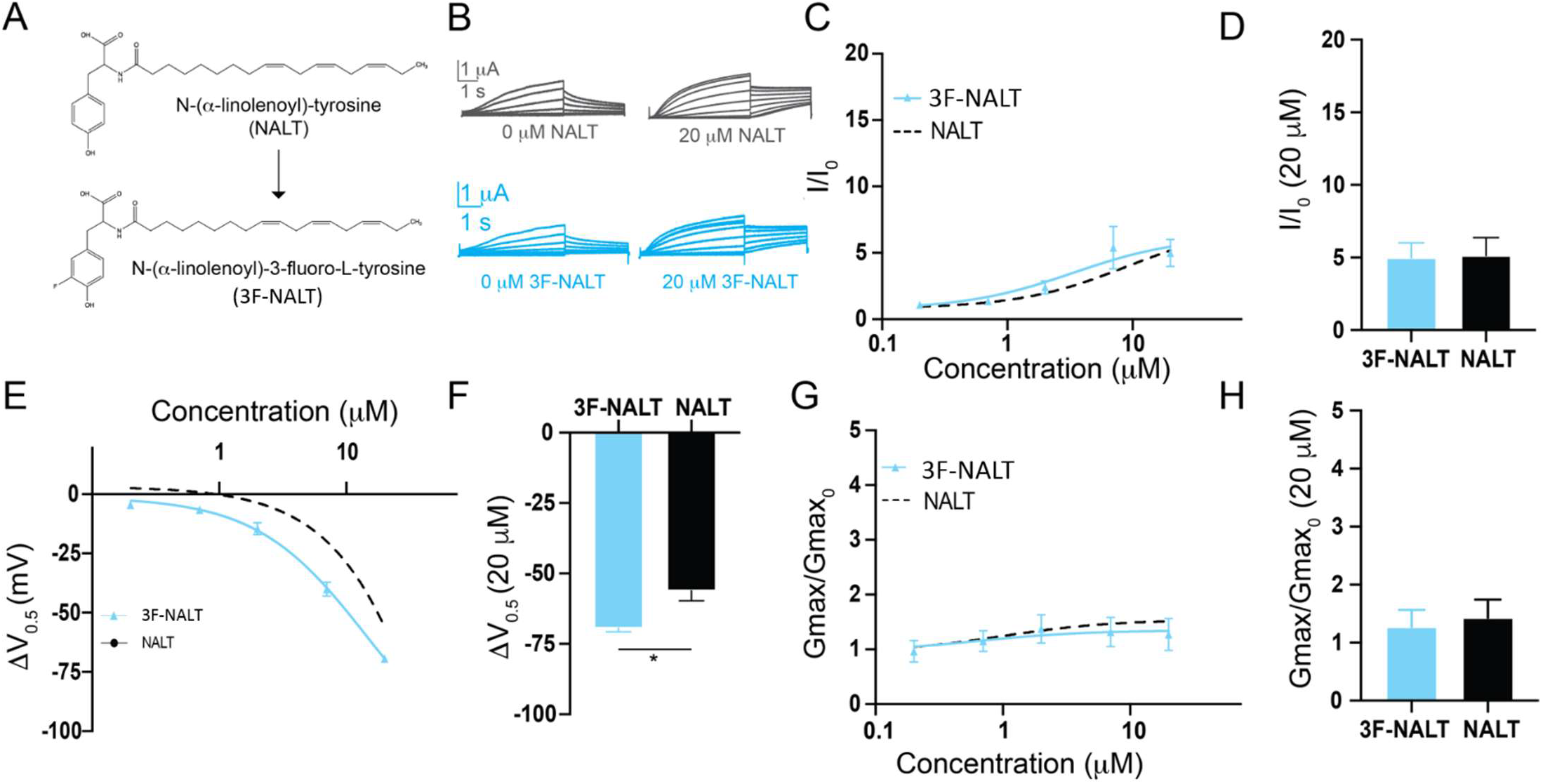
Hydrogen bonding through the distal -OH group of tyrosine PUFAs is important for effects on I_Ks_ channel voltage dependence. **A)** Structures of NALT and 3F-NALT. **B)** Representative traces of NALT (gray) and 3F-NALT (cyan) with 0 μM PUFA (left) and 20 μM PUFA (right). **C**,**E**,**G)** I/I_0_, **E)** ΔV_0.5_, and **G)** G_max_ dose response curves for NALT (black dashed line) (n=4) and 3F-NALT (cyan) (n=3). **D**,**F**,**H)** Maximum effects on **D)** I/I_0_, **F)** ΔV_0.5_, and **H)** G_max_ (at 20 μM) for 3F-NALT (n=3) and NALT (n=4). Values for all compounds and concentrations available in Figure 4-source data 4.

### Aromatic PUFAs appear to activate the I_Ks_ channel in similar mechanisms as non-aromatic PUFAs

To better understand the mechanism of these superior activating aromatic PUFAs we mutated residues previously shown to be important for non-aromatic PUFA activating effects on I_Ks_ channels. The residue R231, located in the voltage sensor (S4) (Fig. 5A), has been previously shown to be important for the V_0.5_ shifting effect of non-aromatic PUFAs^22^. We tested Lin-tyr, the largest V_0.5_ shifting aromatic PUFA, on the I_Ks_ channel with the mutation R231Q+Q234R to assess if R231 is also important for the aromatic PUFA V_0.5_ shifting mechanism. The additional mutation Q234R is necessary to preserve the voltage dependence of activation in I_Ks_ channels with the R231Q mutation^22,28,29^. The V_0.5_ shifting effect of Lin-tyr was significantly decreased from −74.4mV ± 4.1 at 20 μM in the wild-type (WT) I_Ks_ channel to −36.5mV ± 7.3 at 20 μM with the R231Q+Q234R mutation (p = 0.0021**; Fig. 5B-C). This reduction indicates that R231 contributes to more than half of the voltage dependence shifting effect of Lin-tyr. The remaining shift is most likely due to PUFA head group interactions with other nearby S4 charges such as R228 and Q234R.

**Fig. 5.**
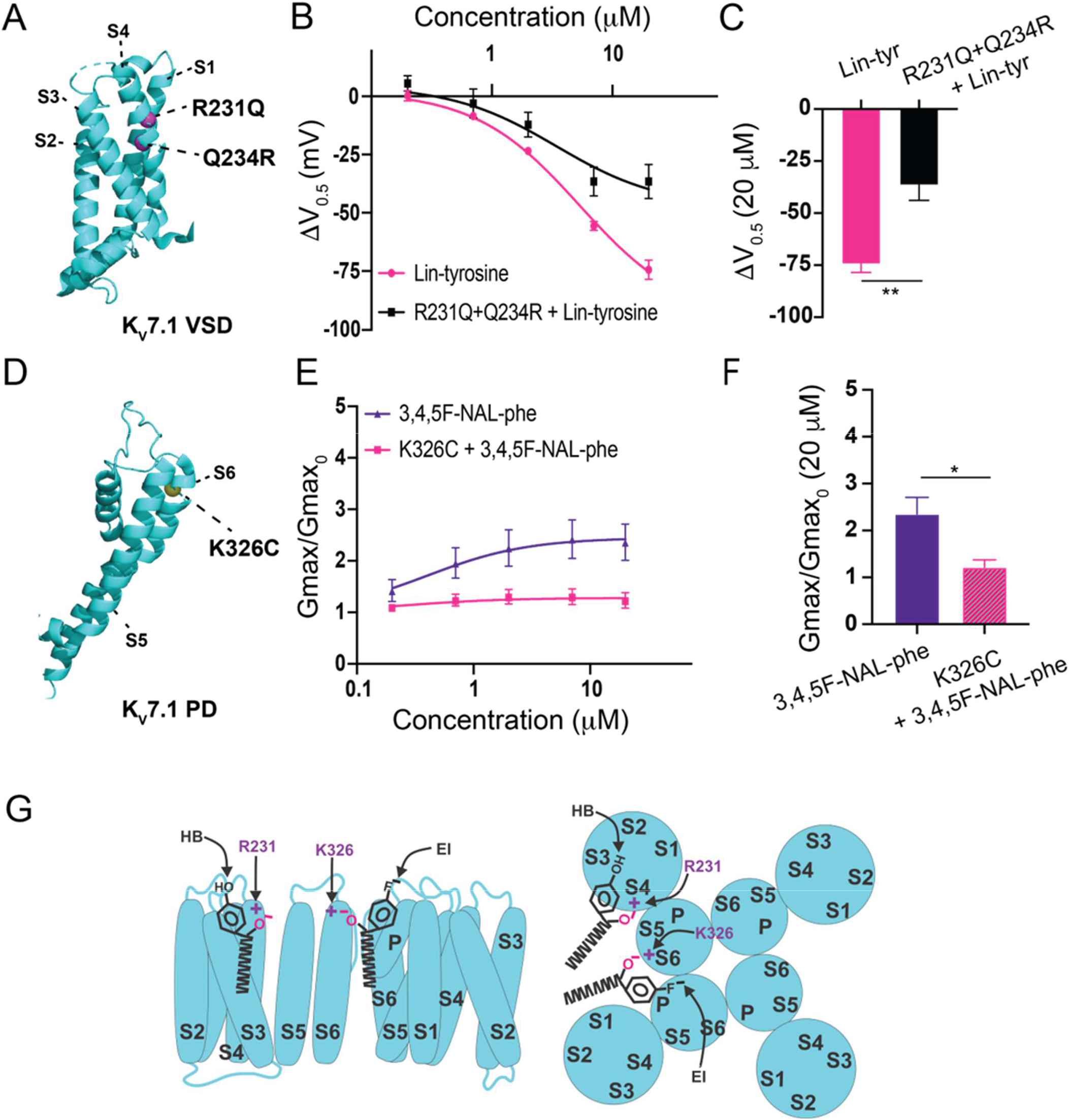
Proposed mechanisms of aromatic PUFAs. **A)** Structure of K_V_7.1 voltage sensing domain (VSD) (based on PDB: 6V00A projected using PyMOL Software (Schrödinger, L. & DeLano, W., 2020. *PyMOL*). Pink spheres indicate mutated residues in the S4 segment, R321Q-Q234R – which are implicated in PUFA-mediated effects on voltage dependent activation. **B)** ΔV_0.5_ dose response curve for WT K_V_7.1/KCNE1 + Lin-tyr (pink) (n=4) and K_V_7.1-R231Q-Q234R/KCNE1 + Lin-tyr (black) (n=4). **C)** Maximum effects on ΔV_0.5_ (at 20 μM) for WT K_V_7.1/KCNE1 + Lin-tyr (n=4) and K_V_7.1-R231Q-Q234R/KCNE1 + Lin-tyr (n=4). **D)** Structure of K_V_7.1 pore domain (PD). Yellow spheres indicates mutated residue in the S6 segment, K326C – which is implicated in PUFA-mediated effects on maximal conductance. **E)** G_max_ dose response curve for WT K_V_7.1/KCNE1 + 3,4,5F-NAL-phe (purple) (n=5) and K_V_7.1-K326C/KCNE1 + 3,4,5F-NAL-phe (pink) (n=3). **F)** Maximum effects on G_max_ (at 20 μM) for WT K_V_7.1/KCNE1 + 3,4,5F-NAL-phe (n=5) and K_V_7.1-K326C/KCNE1 + 3,4,5F-NAL-phe (n=3). **G)** Model for aromatic PUFAs effect on K_V_7.1/KCNE1 channels, side view (left) and top view (right). One site is between S4 and S5: Aromatic PUFAs shift the voltage dependence of opening by stabilizing the upstate of S4 by an electrostatic interactions between R231(+) and the carboxyl group (O^-^) of the PUFA. A hydrogen bond (HB) by the hydroxyl group (OH) at the para site of the aromatic ring of the PUFA stabilize the PUFA in this site. Another site is between S6 and S1: Aromatic PUFAs increase the maximum conductance by an electrostatic interactions between K326(+)and the carboxyl group (O^-^). An electrostatic interaction (EI) by the para fluorine (F^-^) stabilize the PUFA in this site. Values for all compounds and concentrations available in Figure 5-source data 5.

The residue K326, located near the pore, has been previously shown to be important for the G_max_ increasing effect of non-aromatic PUFAs^22^. We tested 3,4,5F NAL-phe, the largest G_max_ increasing aromatic PUFA, on the I_Ks_ channel with the mutation K326C to assess if K326 is also important for the aromatic PUFA G_max_ increasing mechanism (Fig. 5D). The G_max_ increasing effect of 3,4,5F NAL-phe was significantly decreased from 2.4 ± 0.4 at 20 μM in the WT I_Ks_ channel to 1.22 ± 0.2 at 20 μM (p = 0.0287*; Fig. 5E-F). This reduction indicates that K326 is necessary for 3,4,5F NAL-phe’s G_max_ increasing effect. Therefore, aromatic PUFA analogues modulate the I_Ks_ channel via two independent interactions with residues in S4 (R231) and S6 (K326), consistent with the previously described activation mechanisms of PUFAs on I_Ks_ channels (Fig. 5G).

### Residue T224 in the S3-S4 loop is a novel locus for hydrogen bond formation between the I_KS_ channel and tyrosine PUFAs

Our experiments using fluorinated NALT (NAL-3F-tyr) to improve the hydrogen bonding capacity of the tyrosine head group demonstrated that hydrogen bonding by the tyrosine’s para-hydroxyl group is the reason for the large effect of PUFAs with tyrosine head groups on the I_Ks_ channel voltage-dependent activation. To identify the residue with which the tyrosine head group hydrogen bonds, we mutated residues in the S3-S4 loop capable of hydrogen bond formation. We individually mutated serine 217 (S217A), glutamine 220 (Q220L), threonine 224 (T224V), and serine 225 (S225A) and compared the effects of NALT on mutated channels compared to the WT I_Ks_ channel (Fig.6A-B). We found that S217A, Q220L, and S225A showed similar maximum shifts in voltage-dependent activation compared to the wild-type channel (WT + NALT: −56.1 ± 3.6 mV; S217A + NALT: −65.9 ± 3.7 mV; Q220L + NALT: −59.5 ± 11.1 mV; S225A + NALT: −52.4 ± 3.7 mV at 20 μM, ns); Fig. 6C-D). However, the T224V mutation significantly attenuated the leftward shift in the voltage dependence of activation in response to NALT application from −56.1 ± 3.6 mV in WT channels to −32.1 ± 7.0 at 20 μM (p = 0.03*; Fig. 6D). To determine whether this effect was specific to compounds with the ability to form hydrogen bonds we compared the effects of hydrogen-bonding NALT and non-hydrogen-bonding NAL-phe on T224V mutant channels (Fig. 6E). In contrast to the attenuation of the overall voltage shift observed when NALT was applied to the T224V, there was no difference in the voltage-shifting effects of NAL-phe between the T224V mutant and WT channels (WT + NAL-phe: −12.5 ± 3.8 mV; T224V + NAL-phe: −7.8 ± 2.1 mV at 20 μM, ns (Fig. 6F-G). These data demonstrate that the T224V mutation only reduces the efficacy of aromatic PUFAs that contain a hydrogen-bonding group like tyrosine. As a result, we have identified a novel interaction between the S3-S4 loop residue T224 and hydrogen bonding moieties of aromatic PUFA head groups (Fig. 6H).

**Fig. 6.**
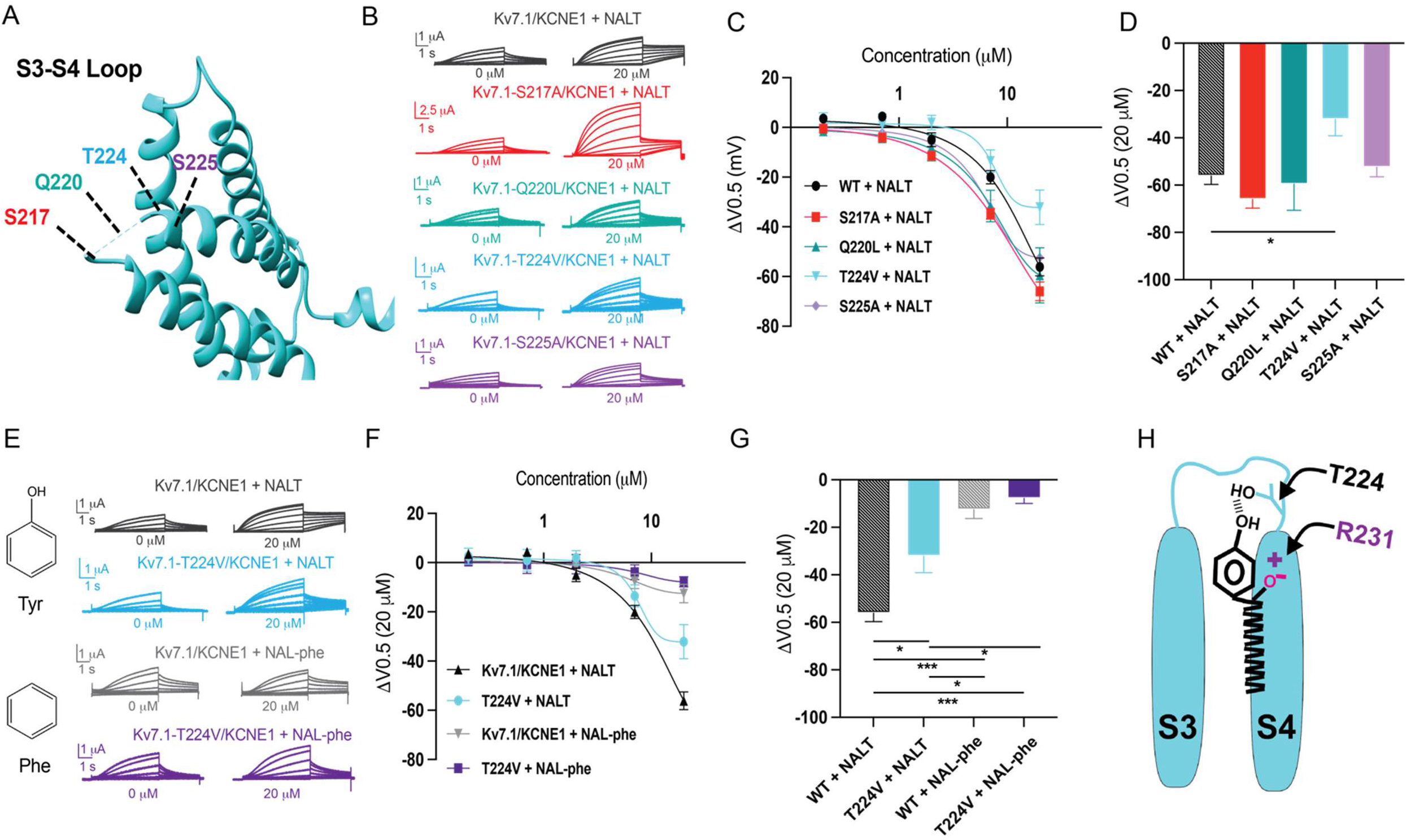
S3-S4 loop is the locus for hydrogen bonding interactions with tyrosine PUFAs. **A)** Top view of K_V_7.1 voltage sensing domain (VSD) highlighting mutated residues in the S3-S4 loop. **B)** Representative traces of WT K_V_7.1/KCNE1 (black), K_V_7.1-S217A/KCNE1 (red), K_V_7.1-Q220L/KCNE1 (teal), K_V_7.1-T224V/KCNE1 (cyan), and K_V_7.1-S225A/KCNE1 (purple) with 0 μM (left) and 20 μM (right) NALT. **C)** ΔV_0.5_ dose response curve for WT K_V_7.1/KCNE1 (n=4), K_V_7.1-S217A/KCNE1 (n=5), K_V_7.1-Q220L/KCNE1 (n=3), K_V_7.1-T224V/KCNE1 (n=4), and K_V_7.1-S225A/KCNE1 (n=7) with NALT. **D)** Maximum effects on ΔV_0.5_ (at 20 μM) for WT and S3-S4 loop mutations (Asterisks indicate statistically significant differences determined by One-way ANOVA). **E)** Representative traces of WT K_V_7.1/KCNE1 with NALT (black) and NAL-phe (gray) compared to K_V_7.1-T224V/KCNE1 with NALT (cyan) and NAL-phe (dark purple), K_V_7.1-S217A/KCNE1 (red), K_V_7.1-Q220L/KCNE1 (teal), K_V_7.1-T224V/KCNE1 (cyan), and Kv7.1-S225A/KCNE1 (purple) with 0 μM (left) and 20 μM (right) NALT **F)** ΔV_0.5_ dose response curve for WT K_V_7.1/KCNE1 and KV7.1-T224V/KCNE1 with NALT and NAL-phe. **G)** Maximum effects on ΔV_0.5_ (at 20 μM) for WT K_V_7.1/KCNE1 (n=4) and K_V_7.1-T224V/KCNE1 with NALT (n=4) and NAL-phe (n=4)(Asterisks indicate statistically significant differences determined by One-way ANOVA). **H)** Model for aromatic PUFAs effect on the voltage dependence of K_V_7.1/KCNE1 channels, illustrating the electrostatic interaction between negatively charged PUFA head groups and R321, in addition to the hydrogen bonding interaction between the para-hydroxyl group of tyrosine PUFAs and T224. Values for all compounds and concentrations available in Figure 6-source data 6.

## Discussion

We have found that PUFA analogues with tyrosine head groups are strong activators of the cardiac I_Ks_ channel. Tyrosine PUFAs shift the voltage dependence of activation to negative potentials and increase the maximal conductance which together contribute to increases in overall I_Ks_ current. The tyrosine head group is an aromatic ring with a distal -OH group in the para-position. Tyrosine PUFA analogues have the potential to interact with the I_Ks_ channel through several candidate mechanisms involving either the aromatic ring or the -OH group (or both). The aromatic ring could modulate I_Ks_ channel function through cation-pi interactions with positively charged groups on the I_Ks_ channel. In addition, the -OH group could participate in electrostatic interactions and/or act as a hydrogen bond donor. In this work, we elucidate the mechanisms of this PUFA-induced activation of the I_Ks_ channel by applying PUFA analogues with modified aromatic head groups designed to test specific chemical interactions between the PUFA head group and the I_Ks_ channel.

If cation-pi interactions were the primary mechanism through which tyrosine PUFAs activate the I_Ks_ channel, we would expect similar activating effects of PUFA analogues with aromatic rings that lack the –OH group, such as phenylalanine. However, PUFA analogues with phenylalanine head groups (Lin-phe and NAL-phe) do not activate the I_Ks_ channel to the same degree as PUFA analogues with a tyrosine head group (Lin-tyr and NALT) and display significant reductions in efficacy for increases in I/I_0_ and shifts in the V_0.5_. Further evidence that cation-pi interactions are not a predominant mechanism for I_Ks_ channel activation by tyrosine PUFA analogues comes from experiments applying fluorinated phenylalanine PUFAs (4F-NAL-phe and 3,4,5F-NAL-phe), which can be used as a tool to probe cation-pi interactions in ion channel function^30^. Pless et al., 2014 demonstrated that tri-fluorination of phenylalanine disperses the electrostatic surface potential which is necessary for cation-pi interactions^30^. Disruption of the electrostatic surface potential through addition of fluorine atoms to the NAL-phe head group (3,4,5F-NAL-phe), therefore, is expected to reduce the efficacy of 3,4,5F-NAL-phe in comparison to NAL-phe alone. However, we find the opposite when we apply 3,4,5F-NAL-phe to the cardiac I_Ks_ channel, and see that 3,4,5F-NAL-phe is a more potent activator of the I_Ks_ channel compared to NAL-phe alone. Together, these data suggest that cation-pi interactions are not the primary mechanism through which these aromatic PUFA analogues activate the cardiac I_Ks_ channel.

When we look at several fluorinated and brominated phenylalanine PUFA analogues, we find specifically that 3,4,5F-NAL-phe has significantly greater effects on I/I_0_ and ΔV_0.5_ compared to NAL-phe alone. While not statistically significant, 4Br-, 4F-, and 3,4,5F-NAL-phe also lead to some of the most consistent increases in G_max_ among the PUFA analogues tested in this work, with each of these compounds leading to a two-fold increase in G_max_. These data suggest that aromatic PUFA analogues with highly electronegative atoms on the distal end of the aromatic head group have the most pronounced effects on the maximal conductance of the I_Ks_ channel. Although, brominated and fluorinated phenylalanine analogues increase the maximal conductance of the I_Ks_ channel, these modified PUFAs still fail to recapitulate the leftward ΔV_0.5_ observed with tyrosine PUFA analogues. While the -OH group of tyrosine PUFA analogues is indeed strongly electronegative, it can also act as a hydrogen bond donor. When we applied a fluorinated tyrosine PUFA (3F-NALT) to increase hydrogen bonding abilities, we found that this leads to a stronger leftward shift in the voltage dependence of I_Ks_ activation. This suggests that hydrogen bonding via the -OH group contributes to the left-shifting effects of voltage dependent activation through effects on the I_Ks_ channel voltage sensor. Most notably, these results suggest that specific modifications to the aromatic PUFA head group can preferentially improve either the voltage-shifting or maximal conductance effects of PUFA analogues. Our data suggests that adding highly electronegative groups to an aromatic ring, such as bromine and fluorine, most consistently improve the maximal conductance increasing effects and reduce voltage dependence shifting effects relative to PUFA analogues with a tyrosine or phenylalanine head group. On the other hand, we found that reducing the pK_a_ of the -OH group (and increasing the potential for hydrogen bonding), while leaving the effect on G_max_ intact, preferentially improves the voltage-shifting effects on the I_Ks_ channel.

Previous work has demonstrated that PUFA analogues have two independent effects on I_Ks_ channel activation. PUFA analogues are known to shift the voltage dependence of activation in the I_Ks_ channel through electrostatic effects on the channel voltage sensor^20,31^. This is mediated by interactions of the negative PUFA head group with the outermost positively charged arginine residues located in the S4 segment^25,31^. Recently, though, a second effect on the I_Ks_ channel pore has been reported to influence the maximal conductance of the I_Ks_ channel^22^. This is mediated through electrostatic interactions between the PUFA head groups and a positively charged lysine residue in the S6 segment – K326_22_. In addition, molecular dynamics (MD) simulations with the Kv7.1 (KCNQ1) channel (the pore-forming domain for the I_Ks_ channel)^21^ identified two separate high occupancy sites for linoleic acid: Site 1 at R228 in the S4 segment, and Site 2 at K326 in the S6 segment^21^. We here show that the superior-activating aromatic PUFAs also act on these sites in S4 and S6. To do this we selected the best V_0.5_ shifting aromatic PUFA (Lin-tyr) to test on the I_Ks_ channel with the S4 mutation R231Q. Additionally, we selected the best G_max_ increasing aromatic PUFA (3,4,5F-NAL-phe) to test on the I_Ks_ channel with S6 mutation K326C. The mutation R231Q decreases the V_0.5_ shifting effect of Lin-tyr by half, indicating that Lin-tyr is shifting the voltage dependence by creating an electrostatic interaction with the positive charges on the voltage sensor. Conversely, the mutation K326C almost completely removed the G_max_ increasing effect of 3,4,5F-NAL-phe. We therefore propose that the increased effects of the aromatic PUFAs, compared to non-aromatic PUFAs, are due to the additional hydrogen bonding in Site 1 and electrostatic interactions in Site 2 to better anchor them in these binding sites to increase their effects (Fig. 5G). As mentioned above, we also show that the aromatic rings have the potential to be modified to give preferential effects on either the I_Ks_ channel voltage sensor or channel pore.

Our experiments with NAL-Phe and 3F-NALT show that the hydrogen bonding capacity of the -OH on the tyrosine of NALT is necessary for it to have superior voltage dependence shifting effect. We further discovered the specific details of the hydrogen bond interactions between this -OH group of NALT and the S3-S4 loop of the I_Ks_ channel. We mutated all residues capable of hydrogen bonding in the S3-S4 loop, removing their ability to hydrogen bond and tested if this changed the NALT voltage dependence shifting effect. The voltage dependence of mutations S217A, Q220L, and S225A was shifted to the same degree by NALT as the wild type I_Ks_ channel. However, the voltage dependence of mutation T224V was shifted significantly less than the WT I_Ks_ channels. This shows that the -OH group on the tyrosine of NALT hydrogen bonds with T224V thereby improving the PUFA’s ability to shift the voltage dependence. This hydrogen bond interaction between PUFAs and the 3-4 loop of the I_Ks_ channel is a novel mechanism to increase the effect of PUFAs to activate the I_Ks_ channel. These data suggest that drugs designed to target this interaction would be more effective at shifting I_Ks_ channel voltage dependence.

Overall, our findings suggest that different aromatic PUFA analogs not only increase PUFA efficacy on activating the I_Ks_ channel, but their specific effects on I_Ks_ function can be modulated independently, either increasing the maximal conductance or voltage-shifting effect. This novel mechanistic understanding of how aromatic PUFAs have these increased effects on the I_Ks_ channel may help to aid drug development for Long QT Syndrome. This data provides insight into how PUFA activation of the I_Ks_ channel can be both increased and tailored to specific I_Ks_ channel deficiencies, such as shifts in voltage dependence and decreases in maximal conductance.

## Acknowledgements

This work was supported by the Swedish Research Council (2021-01885, to S.I.L), the European Research Council (ERC) under the European Union’s Horizon 2020 research and innovation program (grant agreement No. 850622 to S.I.L), and the National Institutes of Health (R01HL131461, to H.P.L.). We thank Jason D Galpin and Christopher A. Ahern (University of Iowa; R24 NS104617-04 The Facility for Atomic Mutagenesis) and Xiongyu Wu (Linkoping University) for synthesis of different aromatic PUFAs.

## Materials Availability Statement

Mutations and newly synthesized PUFAs are available from the corresponding author on reasonable request.

